# Integrated multi-omics analysis of *Klebsiella pneumoniae* revealed the discrepancy in flavonoid effect against strain growth between Rutin and Luteolin

**DOI:** 10.1101/2024.02.20.581226

**Authors:** Zhibin Wang, Wanxia Shen, Yuejiao Li, Xiaoyun Wang, Guofeng Xu, Xiefang Yuan, Hongmei Tang, Ning Ma, Xiaolin Zhong, Xing Wang

## Abstract

The emergence of drug resistant pathogenic bacteria is increasingly challenging conventional antibiotics. Plant derived flavonoids are always considered as potential alternatives to antibiotics due to their antimicrobial properties. However, the molecular mechanisms by which flavonoids inhibit pathogenic microorganisms’ growth are not fully understood. In order to better understand the inhibitory mechanism of flavonoids, two flavonoids were used to incubate *Klebsiella pneumoniae* ATCC700603. After incubation for 4 hours, both the metabolomic and transcriptomic analysis were performed. In present study, 5,483 genes and 882 metabolites were measured. Compared to wild control, the Rutin and Luteolin induced 507 and 374 differentially expressed genes (DEGs), respectively. However, the number of differential abundant metabolites (DAMs) were the same. The correlation between DEGs and DAMs were studied. The top 10 correlated DEGs and DAMs were identified in each comparative groups. Our results showed that, compared to Luteolin, Rutin induced the accumulation of metabolites and suppressed genes’ expression. Our results provided an explanation for the disparate effects of two flavonoids and demonstrated the inhibitory mechanism of Rutin on strain growth.

## 1. Introduction

Infections caused by pathogenic bacteria are threatening human beings’ health worldwide [1]. These bacteria are easily to develop drug resistance during antibiotics treatment processes [2]. Once obtained resistance, bacteria are often difficult to eradicate by antibiotics [3]. In addition, drug resistant bacteria have evolved to spread resistance within populations via horizontal gene transfers, as well as gene mutations, which further lead to persistent infections [4, 5]. As a result, new strategies and approaches in finding effective antibacterial agents that are not easily to generate drug resistance are becoming more and more urgent.

Plant derived flavonoids are bioactive compounds in traditional herbal medicines and have been considered as potential antibacterial agents due to their antimicrobial properties [6]. With a C6-C3-C6 skelecton, flavonoids represent a large group of plant secondary metabolites [7, 8]. These flavonoids are consist of more than 8,000 compounds and provide a huge resource pool for novel antibacterial agents discovery [9]. More importantly, with low toxicity and nearly no side effects, flavonoids are much safer in clinical application [10].

The antibacterial activity of flavonoids has been extensively investigated and well established [11, 12]. Flavonoids extracts derived from *Quercus brantii* L. fruit exhibited significant inhibitory effect on both *Staphylococcus aureus* (MIC 500 µg/mL) and *Enterococcus fecalis* (MIC 600 µg/mL), respectively [13]. The extracts from a shrub (*Sambucus australis*) also showed antibacterial activity against *Salmonella typhimurium* (MIC 250 µg/mL) and *Klebsiella pneumoniae* (MIC 250 µg/mL) [14]. Regarding to a single compound, naringenin showed inhibitory effect on clinical strains of clarithromycin-resistant *Helicobacter pylori*, vancomycin-resistant *E. faecalis*, methicillin-resistant *S. aureus*, and betalactam-resistant *Acinetobacter baumannii* and *K. pneumoniae* [15]. In addition, some other reports also demonstrated that flavonoids, such as naringenin and apigenin showed enhanced antimicrobial effect when used in combination with various conventional antibiotics [16, 17].

Although the antibacterial effects of flavonoids are widely discussed, the inhibitory and/or ineffective mechanism of flavonoids is still not clear. Previous studies uncovered that some known antibacterial mechanism of flavonoids varied [18]. Briefly, the flavonoids can inhibit the cell envelope synthesis [19], bacterial motility [20], nucleic acid synthesis [21], biofilm formation [22] and quorum sensing [23] in various pathogenic bacteria. Besides, some flavonoids showed up to sixfold stronger antibacterial activities than conventional antibiotics [12]. Nevertheless, these plant derived components remain a poorly characterized reservoir of anti-infective agents [24].

In our previous studies, we evaluated the antioxidant and antibacterial effects of 10 flavonoids. Nevertheless, consistent with some other studies [25, 26], we found most flavonoids were ineffective in inhibiting the growth of *K. pneumoniae* except Rutin [27]. Most previous antibacterial studies performed the paper disk [28] and/or minimum inhibition concentration (MIC) evaluation [29] and confirmed antibacterial effects of some flavonoids. To reveal the inhibitory mechanism of Rutin and uncover the reason that the activity of two flavonoids varied, we used omics methods in present study. Rutin (effective) and Luteolin (ineffective) were used to incubate *K. pneumoniae* bacterium, respectively. After incubation, bacteria cells were collected for both RNA-seq and metabolome sequencing. The relationship between differential abundant metabolites (DAMs) and differentially expressed genes (DEGs) was assessed. The integrated analysis provide fresh insights into the inhibitory effects of flavonoids on *K. pneumoniae* growth.

## 2. Materials and methods

### 2.1 Bacteria growth conditions and treatments

The standard bacterial strain *K. pneumoniae* ATCC700603 was purchased from China National Health Inspection Center. This strain was cultured in Luria-Bertani broth (LB, Haibo, Qingdao, China) at 37°C. Rutin and Luteolin were purchased from Aladdin Chemistry Co., Ltd. (Shanghai, China). The flavonoids stocks were prepared in mixture of ethanol and methyl sulfoxide (DMSO) and the concentration was made to 1024 µg/mL. All chemicals used in this study were of chromatography grade. Ultra pure water was used throughout all experiments.

Briefly, 10mL bacterial suspension (10^8 cfu/mL) was transferred to a flask of 89 mL LB liquid culture. Then, 1mL of each dissolved flavonoid stock was added to each flask, respectively, making the final volume to 100 mL in each flask. All experiments were performed in triplicate. The empty control was conducted in 3 flasks filled with 90 mL LB and 10 mL bacterial cells. The flasks were covered and incubated for 4h at 37°C with shaking (150 rpm). Then, two specimens were randomly selected from each treatment for further study.

### 2.2 RNA extraction, library preparation and illumina Hiseq sequencing

The bacterium cells were collected from 10 mL culture medium after centrifugation at 1000 × g for 10 min at 4°C. Total RNA was isolated from bacterial cells using TRIzol® Reagent (Invitrogen, Carlsbad, CA, USA), according to the method [30]. RNA quality was determined using 2100 Bioanalyser (Agilent, CA, USA) and quantified by the NanoDrop ND-2000 spectrophotometer (NanoDrop Technologies, Delaware, USA). High-quality RNA sample (OD260:280=1.8∼2.2, OD260:230≥2.0, RIN≥6.5, 23S:16S≥1.0) was used to construct library.

Briefly, a total amount of 5 μg RNA per sample was used for library construction. Firstly, the mRNA was interrupted by ion using TruseqTM Stranded RNA sample prep Kit (Illumina, San Diego, USA). Then the dUTP was used instead of dTTP in the second chain synthesis and the index adaptor was added after double strained cDNA synthesis using TruseqTM Stranded RNA sample prep Kit. After that, the second chain of cDNA was degraded using UNG (Uracil-N-Glycosylase) enzyme (Illumina, San Diego, USA). The library was enriched by PCR (polymerase chain reaction) using cBot Truseq PE Cluster Kit v3-cBot-HS (Illumina, San Diego, USA). Secondly, the library was separated and purified using 2% certified low range ultra agarose gel electrophoresis (Bio-Rad, Herales, USA). The target gene fragments were isolated by gel extraction kit. Then, the library was amplified using a cBot Truseq PE Cluster Kit v3-cBot-HS (Illumina, San Diego, USA). At last, the library preparations (quantified by TBS380, paired-end libraries) were sequenced by Illumina NovaSeq 6000 platform (150 bp*2, Beijing iGeneTech Biotechnology Co., Ltd., Beijing, China). The raw paired end reads were trimmed and quality controlled by Trimmomatic with parameters (SLIDINGWINDOW:4:15 MINLEN:75) (version 0.36). Then clean reads were separately aligned to reference genome with orientation mode using Rockhopper software (http://cs.wellesley.edu/btjaden/Rockhopper/).

The reference genome (GenBank assembly accession: GCA_011045595.1) was extracted from Genome Database for *Klebsiella pneumoniae* (BioProject: PRJNA559783). The index of the reference genome was built with Bowtie 2 (2.2.6), and the clean reads were mapped to the reference genome using Tophat (2.0.14). HTSeq (0.9.1) was applied to the expression and quantification levels of genes to calculate reads per kilobase of transcript per million reads mapped (RPKM). To identify DEGs between two compared groups, the expression level for each transcript was calculated using the RPKM method. The edgeR (https://bioconductor.org/packages/release/bioc/html/edgeR.html) was used for differential expression analysis. The DEGs between two samples were selected using the following criteria: the logarithmic of fold change was greater than 2 and the false discovery rate (FDR) should be less than 0.05. To understand the functions of the differentially expressed genes, GO (Gene Ontology) functional enrichment and KEGG [59, 60] pathway analysis were carried out by Goatools (https://github.com/tanghaibao/Goatools) and KOBAS (http://kobas.cbi.pku.edu.cn/home.do) respectively.

### 2.3 Metabolites measurement and analysis

The freeze-dried samples were mashed with liquid nitrogen, and 50 mg of powder for each sample was transferred into a 1.5 mL EP tube containing 5 mL precooled methanol/water (V:V=3:1) and subjected to vigorous vortexing. Then, the samples were centrifuged at 13,000 rpm for 15 min at 4 °C and the filtered supernatant (0.22 μm membrane filter) was blow-dried with nitrogen. The samples were redissolved with 50 μL isopropanol/acetonitrile/water (V:V:V=2:7:1) and extracted by centrifugation before LC-MS/MS analysis. The extracts were separated on the UPLC system (Dionex UltiMate Ultimate 3000) equipped with ACQUITY UPLC HSS T3 column (100 mm*2.1 mm, 1.7 μm, Waters, USA). The conditions were as follows: column temperature, 40 °C injection volume, 1.5 μL; flow rate, 0.35 mL/min. The mobile phases were water (0.1% formic acid) (phase A) and acetonitrile (phase B). The metabolites were eluted with the following gradient: 0∼1.0 min, 5% B; 1.0∼9.0 min, 5∼100% B; 9.0∼12.0 min, 1000% B; 12.0∼15.0 min 5% B. Samples were inserted into quality control (QC) samples in queue mode to monitor and evaluate the stability of the system and the reliability of the experimental data. Each sample was operated in both positive and negative ion modes by the electrospray ionization (ESI) source parameters. Mass spectrometry was carried out by the Q-Exactive spectrometer (Thermo Scientific, San Jose, CA, USA) after the sample was separated by UPLC. The MS conditions were as follows: mass range (m/z) 80∼1200; aux gas flow rate, 15 arb; sheath gas flow rate, 45 arb; source temperature, 320 ° C; spray voltage, 3.5 kV in positive mode and

3.2 kV in negative mode, respectively. After the original data were adjusted for peak alignment, peak area extraction, retention time correction and feature extraction using the Compound Discoverer 3.0 program, the metabolite structure was identified by accurate mass MS matching (<25 ppm), and the MS1 and MS2 matching search MZcloud and ChemSpider database were used to determine the metabolite structure. The relative content of metabolites was expressed by peak area. After preprocessed by pattern recognition and Pareto-scaling, the peak area data for all metabolites were further analyzed by SIMCA-P 14.1 software (Umetrics, Umea, Sweden), including unsupervised principal component analysis (PCA). Significantly accumulated metabolites between groups were determined by VIP≥1 and log2FC (fold-change) ≥1.

### 2.4 Gene expression verification

To validate the RNA-seq results, 10 genes were randomly selected for quantitative real-time PCR (qRT-PCR). Primers were designed based on cDNA sequences. The 16S rRNA [31] was used as reference gene. Relative gene expression of each gene was calculated by using 2^-ΔΔCT^method [32]. The fold change in gene expression was obtained by the normalization of each gene to the 16S rRNA internal control.

The qRT-PCR was performed by the 7500 RT-PCR system (Applied Biosystems, Foster City, USA) using a SYBR Green RT-PCR Kit (Qiagen, Hilder, Germany). According to the manufacturer’s instructions, 20 µL PCR reaction mixture were composed of 2 µL of cDNA, 1.5 µl of each pair of target primers (200 nM), 5 µL of ddH_2_O and 10 µL of 2 × SYBR® Green Supermix (Bio-Rad Laboratories, Shanghai, China). The PCR conditions were as follows: 95 °C for 5 min (initial denaturation), followed by 40 cycles of denaturation at 95 °C for 15 s, annealing at 60 °C for 25 s, and extension at 72 °C for 30 s. The specificity of each PCR reaction was confirmed by melting curve analysis.

### 2.5 Statistical analysis

All measurements and calculations are presented as mean of three standard error of the mean. SPSS (V19.0) software and GraphPad Prism (V9.1.0) software were used to conduct data analysis and generate graphs. Student’s t-test was used to compare data between each two groups. Pearson correlation analysis was performed in correlation analysis. A P value < 0.05 was considered as statistically significant correlation for all tests. Cytoscape (V3.8.2) software was used to construct the correlation network.

## 3. Results

### 3.1 Inhibitory activities of Rutin and Luteolin on the growth of K. pneumoniae strain

After overnight incubation on agarose solid medium, bacterial colonies grown at different concentrations of Rutin were inhibited obviously (Fig. 1 A). Moreover, Rutin at higher concentration showed stronger inhibitory activity. However, Luteolin showed no obvious suppression (Fig. 1 B).

**Fig. 1.**
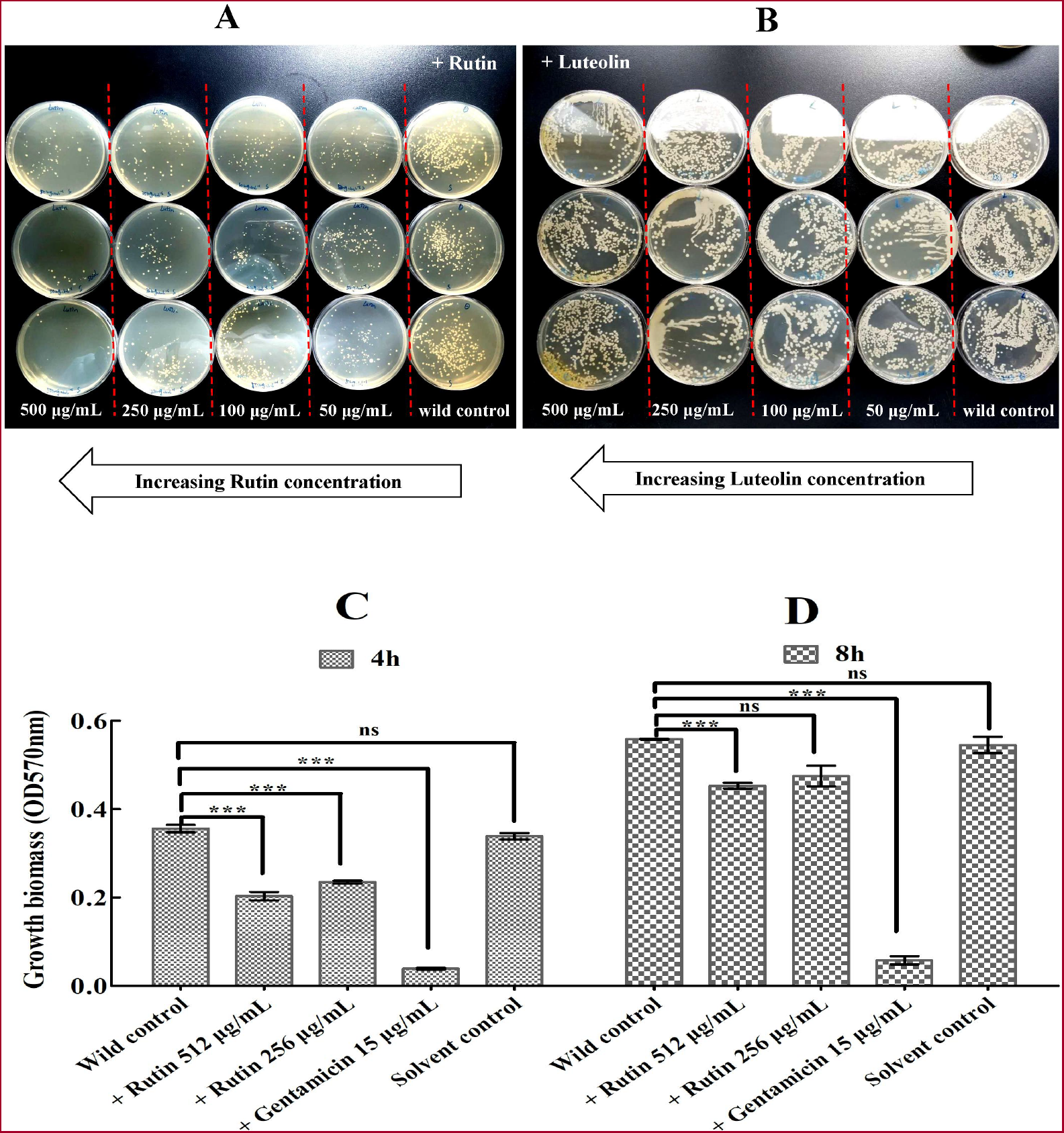
Inhibitory activities of flavonoids against the growth of *K. pneumoniae* ATCC700603. (A) Growth status at different concentrations of Rutin. (B) Growth status at different concentrations of Luteolin. (C) Biomass accumulation at 4h with or without Rutin treatments. (D) Biomass accumulation at 8h with or without Rutin treatments. The wild control was compared to all the other groups, respectively. *** indicates P<0.001 and ns indicates not significant. Gentamicin (15 μg/mL) was used as reference antibiotic. The solvent control showed no significant influence on colony number.

To compare the inhibitory effects of Rutin and Luteolin at different time points, the cut-off OD value were measured at 4 h and 8 h, respectively. Rutin showed inhibitory activity at both 256 μg/mL and 512 μg/mL. However, Luteolin had no obvious inhibition (Fig. 1 C and D).

### 3.2 Differentially expressed genes and metabolites in response to Rutin and Luteolin treatments

An average of 17,066,492 clean reads were obtained per sample from 37,497,710 raw reads for all samples (Supplementary material, Data sheet 1, Table S 1). The Q20 value and Q30 value were 98.96% and 96.23%, respectively. The average mapping rate for all clean reads to the reference transcripts achieved to 65.59%. The raw data of RNA-seq was deposited in NCBI and the reference BioSample accession was SAMN34237808.

The expression levels of the transcripts were calculated utilizing the RPKM approach. Herein, we categorized genes into high, medium and low groups according to their average RPKM value for distinction. In all 6 libraries, more genes were weakly expressed (RPKM ≤1, avg. 2,843), fewer genes had medium (RPKM 1∼10, avg. 1,274) and high expression (RPKM ≥10, avg. 1,366) (Fig. 2 A). Furthermore, genes at a same RPKM level were also compared. The results were shown in venn diagrams (Fig. 2 B∼E). However, there was no big differences in the total amount of genes induced by treatments or without treatments within a same level RPKM group.

**Fig. 2.**
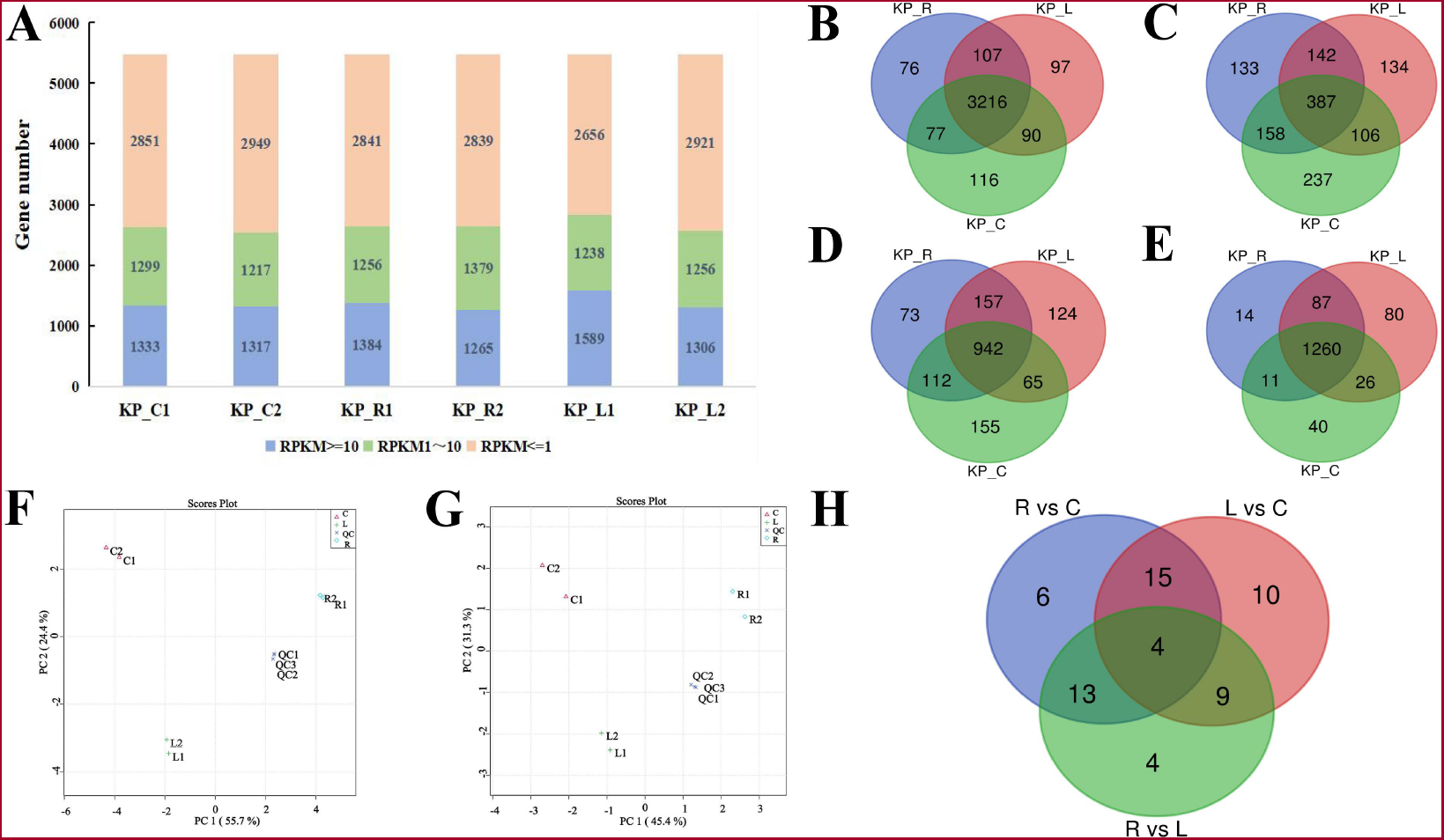
Quantification of transcript expression levels and metabolites. (A) Number of transcripts and their expression levels measured in 6 libraries. (B) Venn diagram showing numbers of all transcripts measured in 3 groups. (C) Venn diagram showing numbers of transcripts at low expression level. (D) Venn diagram showing numbers of transcripts at medium expression level. (E) Venn diagram showing numbers of transcripts at high expression level. (F) Scattered point plots are obtained from PCA analysis in positive ion mode. (G) Scattered point plots are obtained from PCA analysis in negative ion mode. (H) Venn diagram showing numbers of DAMs. C: wild control; R: Rutin treatment (512 μg/mL); L: Luteolin treatment (512 μg/mL). The transcripts shown in venn diagrams were compared in pairs among KP_R group, KP_L group and KP_C group, respectively.

To gain a holistic understanding of the results, pairwise comparisons between wild control group (KP_C), Rutin treatment group (KP_R) and Luteolin treatment group (KP_L) were compared, respectively. However, more DEGs (507) were identified in Luteolin group while less DEGs (374) yielded in Rutin group (Supplementary material, Data sheet 1, Table S 2, 3). These results showed the different influence of Rutin and Luteolin. Further analysis identified 159 DEGs in KP_L vs KP_R (Supplementary material, Data sheet 1, Table S 4).

In metabolomic sequencing, 543 chemical substances under positive ion mode and 339 under negative ion mode were measured using an UPLC-MS detection platform (Supplementary material, Data sheet 2, Table S 1, 2). Under positive ion mode, the principal component 1 (PC1) and principal component 2 (PC2) were accounted for 55.7% and 24.4% of the total variations, respectively (Fig. 2 F). Under the negative ion mode, the PC1 and PC2 were 45.4% and 31.3%, respectively (Fig. 2 G).

In this project, the DAMs were screened according to the VIP (Variable Importance in the Projection) value which were derived from the PLS-DA model analysis. Interestingly, compared to wild control, both Rutin and Luteolin induced 38 DAMs (Fig. 2 H).

### 3.3 Annotation and enrichment analysis of DEGs

The KEGG (Kyoto Encyclopedia of Genes and Genomes) enrichment of KP_R vs KP_C yielded 82 pathways, including metabolic pathways, ABC transporters, microbial metabolism in diverse environments etc (Supplementary material, Data sheet 1, Table S 5). These 82 pathways were mainly classified into 4 categories, which were metabolism, environmental information processing, cellular processes and genetic information processing. Herein, the top 20 pathways which were ranked from most to least basing on the number of DEGs that they contained were exhibited (Fig. 3 A). Similarly, the enrichment analysis of KP_L vs KP_C yielded 95 pathways (Supplementary material, Data sheet 1, Table S 6). Among these pathways, the top 20 enriched pathways were shown in Fig. 3 B. Obviously, among those 20 pathways, 5 (Lipopolysaccharide biosynthesis, Butanoate metabolism, Nitrogen metabolism, 2-Oxocarboxylic acid metabolism and Degradation of aromatic compounds) were unique to Rutin, while another 5 (Benzoate degradation, Pentose phosphate pathway, Phosphotransferase system, Glyoxylate and dicarboxylate metabolism and Purine metabolism) were unique to Luteolin. These findings imply that these two flavonoids exhibited disparate functionalities.

**Fig. 3.**
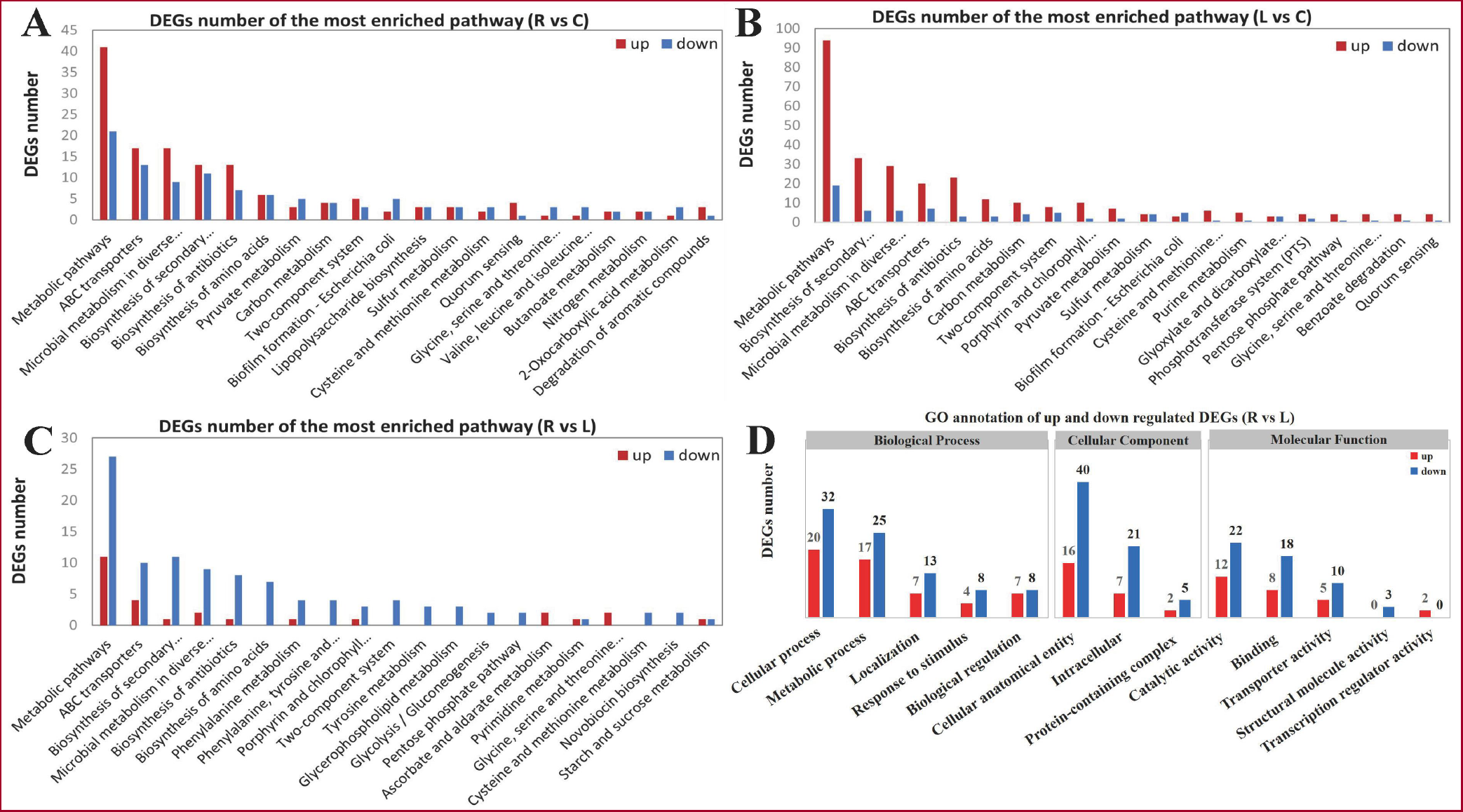
KEGG and GO enrichment of DEGs induced by Rutin and Luteolin treatments. (A) KEGG enrichment of DEGs induced by Rutin, compared to wild control. (B) KEGG enrichment of DEGs induced by Luteolin, compared to control. (C) KEGG enrichment of DEGs derived from KP_L vs KP_R. (D) GO enrichment of DEGs derived from KP_L vs KP_R.

The DEGs from KP_R vs KP_L were also enriched (Supplementary material, Data sheet 1, Table S 7). Among those 57 pathways, 50 were metabolism related pathways, 3 were environmental information processing related pathways, 2 were genetic information processing related pathways and 2 were cellular processes related pathways. The top 20 enriched pathways were represented in Fig. 3 C. Basically, the amount of down regulated DEGs in each pathway was more than that of up regulated ones. More down regulating gene’s number suggested that Rutin could inhibit genes’ expression. At last, the GO analysis reconfirmed this assumption (Fig. 3 D).

### 3.4 Annotation and enrichment analysis of DAMs

Based on the OPLS-DA results, 38 and 38 metabolites were identified as DAMs in KP_R vs KP_C and KP_L vs KP_C, respectively (Supplementary material, Data sheet 2, Table S 3, 4). Further analysis showed that 17 DAMs were found in both comparative groups. However, among those DAMs, 20 were successfully annotated to KEGG in KP_R vs KP_C while only 14 were successfully annotated in KP_L vs KP_C. Besides, 30 DAMs were identified in KP_R vs KP_L (Supplementary material, Data sheet 2, Table S 5).

Obviously, all the DAMs induced by Rutin were up regulated while those induced by Luteolin were both up regulated and down regulated. The DAMs from KP_R vs KP_C and KP_L vs KP_C were enriched into 27 and 37 pathways, respectively (Fig. 4). These pathways were mainly related to metabolism, genetic information processing, environmental information processing and organismal systems. Similarly, the DAMs from the KP_R vs KP_L were also annotated (Fig. 4 C). In this comparative group, both up regulated and down regulated DAMs were found. Together with 7 metabolism related pathways and 2 environmental information processing related pathways, those DAMs were annotated to 9 pathways. The dissimilarity in metabolite profiles induced by two flavonoids were depicted on the correlation heat map (Fig. 4 D). Clearly, the administration of Rutin leaded to the accumulation of a majority of metabolites, while Luteolin exerted inhibitory effects on these metabolites.

**Fig. 4.**
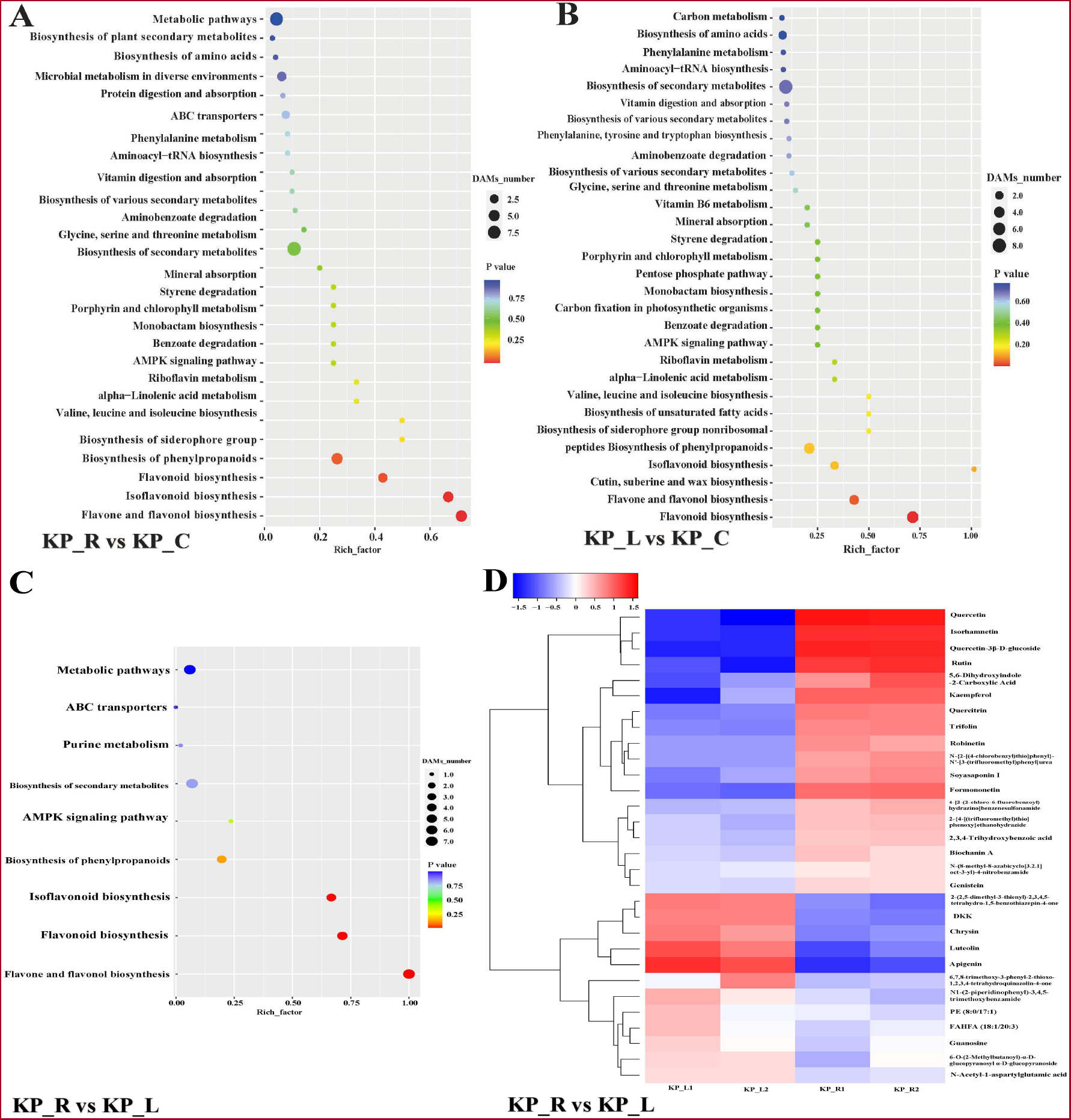
KEGG enrichment and correlation heat map of DAMs. (A) KEGG enrichment of DAMs derived from KP_R vs KP_C. (B) KEGG enrichment of DAMs derived from KP_L vs KP_C. (C) KEGG enrichment of DAMs derived from KP_R vs KP_L. (D) Correlation heat map of DAMs derived from KP_R vs KP_L. For A, B and C, the horizontal axis represents the enrichment factor, that is, the ratio of the number of DAMs enriched in a pathway to the background metabolites. The ordinate represents the function enriched by KEGG. For D, the horizontal and vertical coordinates in the figure represent the different metabolites in comparative group, and the color blocks at different positions represent the correlation coefficients between metabolites, with red representing positive correlation and blue representing negative correlation.

### 3.5 Integrated analysis of transcriptomics and metabolomics

To refine functional pathways, we conducted a comprehensive analysis integrating transcriptomic and metabolomic data. The pathways that were enriched in both the transcriptome and metabolome were screened for further study. From KP_ R vs KP_C, 12 pathways were filtered out (Table 1). Apparently, the administration of Rutin treatment resulted in alterations in 12 pathways, with the predominant involvement of Metabolic pathways (ko01100), ABC transporters (ko02010), and Microbial metabolism in diverse environments (ko01120) which contained a higher number of associated genes. From KP_R vs KP_L, 4 pathways were identified (Table 2). Herein, except for ko00230 pathway (Purine metabolism), all the other pathways can be found in KP_R vs KP_C.

**Table 1.**
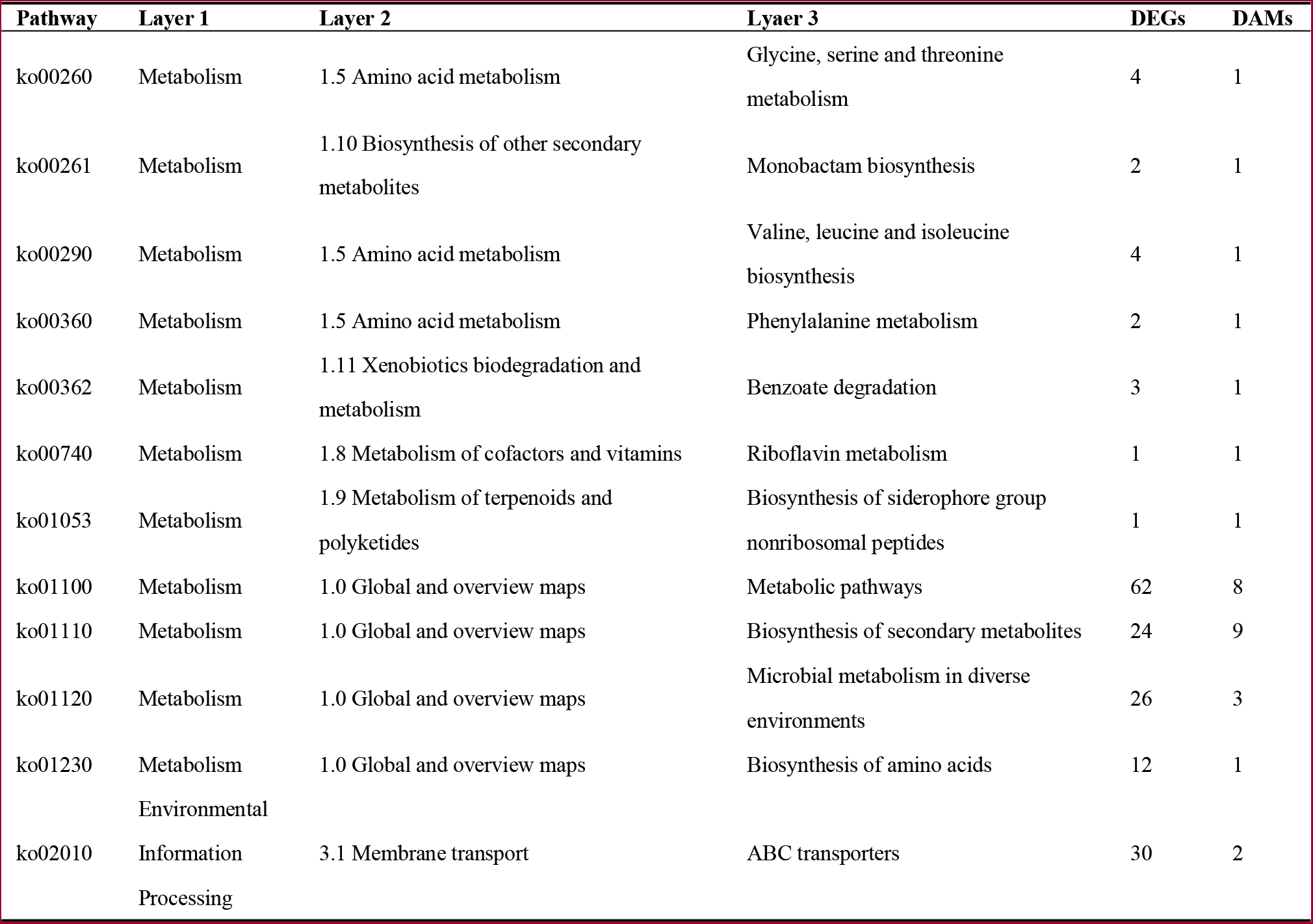
The shared pathways derived from the annotations of both transcriptomic and metabolomic analysis in KP_R vs KP_C.

**Table 2.** The shared pathways derived from the annotations of both transcriptomic and metabolomic analysis in KP_R vs KP_L.

| Pathway | Layer 1 | Layer 2 | Layer 3 | DEGs | DAMs |
| --- | --- | --- | --- | --- | --- |
| ko00230 | Metabolism | 1.4 Nucleotide metabolism | Purine metabolism | 1 | 1 |
| ko01100 | Metabolism | 1.0 Global and overview maps | Metabolic pathways | 38 | 7 |
| ko01110 | Metabolism | 1.0 Global and overview maps | Biosynthesis of secondary metabolites | 12 | 7 |
| ko02010 | Environmental Information Processing | 3.1 Membrane transport | ABC transporters | 14 | 1 |

The correlation between the DEGs and the DAMs were calculated using the Pearson index and represented in heat maps (Supplementary material, Data sheet 3, Figure S1, 2). Only those DEGs and DAMs with significant correlation coefficients were used to build correlation network. The raw data was described in Supplementary material, Data sheet 4 and 5, respectively.

In KP_R vs KP_C, 4 pathways consisted of 23 genes and 9 metabolites were identified (Fig. 5 A). Among these metabolites, C00389 (Quercetin, C15H10O7) and C00188 (L-threonine, C4H9NO3) were the most and least abundant substances, respectively. The amino acid L-threonine, however, was associated with a significantly larger number of genes, including 2 negatively correlated and 14 positively correlated ones. Another vital substance was C00255 (Vitamin B2, C17H20N4O6), which was found to be correlated with 14 genes and simultaneously associated with 3 pathways. Of these 14 genes, 9 were involved in ABC transporters.

**Fig. 5.**
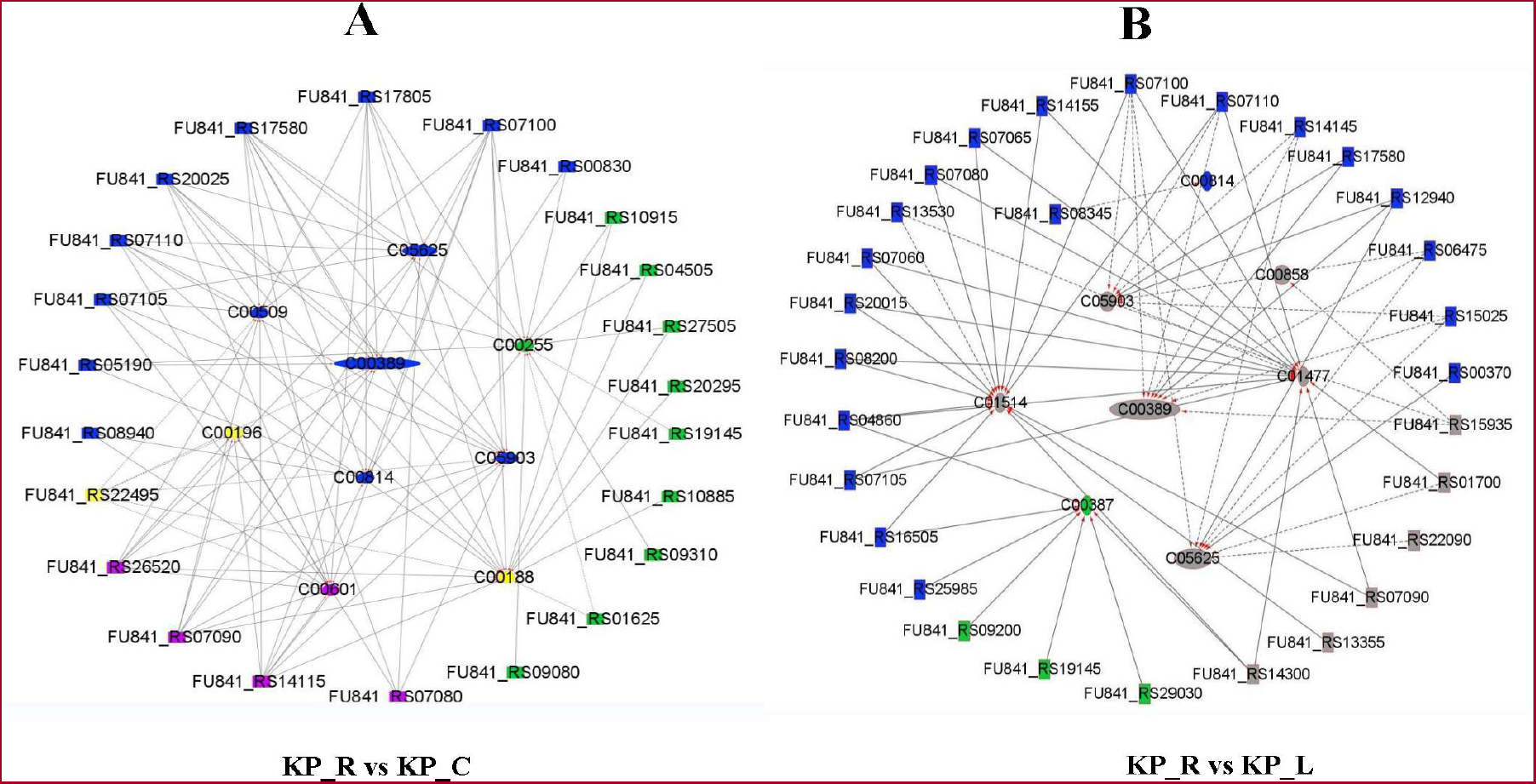
Correlation network of DEGs and DAMs. (A) Correlation network of DEGs and DAMs in KP_R vs KP_C. (B) Correlation network of DEGs and DEMs in KP_R vs KP_L. Blue colour represents metabolic pathways (ko01100), purple colour represents microbial metabolism in diverse environments (ko01120), yellow colour represents biosynthesis of amino acids (ko01230), green colour represents ABC transporters (ko02010), and gray colour represents biosynthesis of secondary metabolites (ko01110). DAMs or DEGs with a same colour were involved in a same pathway. Rectangle represents gene and ellipse represents metabolite. Dash line represents negative correlation and solid line represents positive correlation, respectively.

In KP_R vs KP_L, more DEGs were found to be clustered in the metabolic pathways while less DEGs were found in the ABC transporters pathway (Fig. 5 B). The metabolic pathway contained 20 genes and 1 metabolite while the ABC transporters pathway covered 3 genes and 1 metabolite.The most abundant metabolite was also Quercetin, followed by C05625 (Rutin, C27H30O16), C00858 (Formononetin, C16H12O4), Kaempferol, C00814 (Biochanin A, C16H12O5), C00387 (Guanosine, C10H13N5O5), C01514 (Luteolin, C15H10O6) and C01477 (Apigenin, C15H10O5). The subsequent analysis showed that there were 4 common metabolites observed in both comparative groups. The table (Supplementary material, Data sheet 2, Figure S 6) displays the metabolites that are shared by both comparative groups as well as the specific metabolites. Interestingly, metabolites specific to Rutin treatment were increased, while those specific to Luteolin treatment were decreased.

The top 10 correlated genes and metabolites were listed in Table 3 And Table 4, respectively. In KP_R vs KP_C, the Aspartate kinasegene gene (*thrA*) was considered as a vital gene due to its association with 6 metabolites in Table 3, followed by malonyl-ACP O-methyltransferase (*bioC*) which was associated with 4 metabolites. Gene *thrA* was negatively correlated with 6 metabolites, four of which were specifically produced by Rutin induction. It was reported that L-threonine was one of the subsequent metabolisms in the biosynthetic pathway from L-aspartate to L-homoserine, in which Aspartate kinase (EC 2.7.2.4) was involved [33, 34]. The accumulation of L-threonine was observed and the expression of *thrA* gene was inhibited upon Rutin treatment, suggesting a potential role for Rutin in this processes. In addition, the activity of Aspartate kinase could be inhibited by L-threonine [34]. In present study, the negative correlation between *thrA* and L-threonine coincided with the previous study. As a result, we hypothesized that the other metabolites, like Naringenin and Quercetin played a similar role as L-threonine. Biotin plays an essential role in bacteria growth and *bioC* was reported to encode BioC protein which was the initiator of the synthetic pathway that forms the pimeloyl moiety of biotin [35, 36]. In Table 3, L-threonine and 3 flavonoids showed positive correlation with *bioC* gene. These findings implied that the growth suppression of Rutin may also be associated with biotin biosynthesis.

**Table 3.** The most correlated DEGs and DAMs derived from the correlation network of KP_R vs KP_C group.

| Gene ID | Metabolite ID | Compound | Correlation coefficient | P value | Function |
| --- | --- | --- | --- | --- | --- |
| FU841_RS22495 | C00509 | Naringenin | -0.9660 | 0.0340 | *thrA*; bifunctional aspartate kinase/homoserine dehydrogenase I |
| FU841_RS22495 | C00389 | Quercetin | -0.9610 | 0.0390 | *thrA*; bifunctional aspartate kinase/homoserine dehydrogenase I |
| FU841_RS22495 | C05903 | Kaempferol | -0.9580 | 0.0420 | *thrA*; bifunctional aspartate kinase/homoserine dehydrogenase I |
| FU841_RS22495 | C00188 | L-threonine | -0.9530 | 0.0470 | *thrA*; bifunctional aspartate kinase/homoserine dehydrogenase I |
| FU841_RS22495 | C00196 | 2,3-Dihydroxy benzoic acid | -0.9530 | 0.0470 | *thrA*; bifunctional aspartate kinase/homoserine dehydrogenase I |
| FU841_RS22495 | C00601 | Phenylacetaldehyde | -0.9800 | 0.0200 | *thrA*; bifunctional aspartate kinase/homoserine dehydrogenase I |
| FU841_RS01625 | C00255 | Vitamin B2 | -0.9970 | 0.0030 | *livF*; high-affinity branched-chain amino acid ABC transporter ATP-binding protein LivF |
| FU841_RS01625 | C00188 | L-threonine | -0.9760 | 0.0240 | *livF*; high-affinity branched-chain amino acid ABC transporter ATP-binding protein LivF |
| FU841_RS19145 | C00255 | Vitamin B2 | -0.9720 | 0.0280 | sugar ABC transporter ATP-binding protein |
| FU841_RS07100 | C00389 | Quercetin | 0.9980 | 0.0020 | *eutA*; ethanolamine ammonia-lyase reactivating factor EutA |
| FU841_RS07110 | C00509 | Naringenin | 0.9970 | 0.0030 | *eutC*; ethanolamine ammonia-lyase subunit EutC |
| *FU841_RS07110* | C00188 | L-threonine | 1.0000 | 0.0004 | *eutC*; ethanolamine ammonia-lyase subunit EutC |
| *FU841_RS07090* | C05903 | Kaempferol | 0.9990 | 0.0010 | *eutG*; ethanolamine utilization ethanol dehydrogenase EutG |
| *FU841_RS07090* | C00196 | 2,3-Dihydroxy<br>benzoic acid | 0.9980 | 0.0020 | *eutG*; ethanolamine utilization ethanol dehydrogenase EutG |
| *FU841_RS17580* | C05903 | Kaempferol | 0.9980 | 0.0020 | *bioC*; malonyl-ACP O-methyltransferase BioC |
| *FU841_RS17580* | C00389 | Quercetin | 0.9990 | 0.0010 | *bioC*; malonyl-ACP O-methyltransferase BioC |
| *FU841_RS17580* | C00509 | Naringenin | 0.9990 | 0.0010 | *bioC*; malonyl-ACP O-methyltransferase BioC |
| *FU841_RS17580* | C00188 | L-threonine | 1.0000 | 0.0004 | *bioC*; malonyl-ACP O-methyltransferase BioC |
| *FU841_RS10915* | C00255 | Vitamin B2 | 0.9980 | 0.0020 | *ccmA*; cytochrome c biogenesis heme-transporting ATPase CcmA |

**Table 4.** The most correlated DEGs and DAMs derived from the correlation network of KP_R vs KP_L.

| Gene ID | Metabolite ID | Compound | Correlation coefficient | P value | Function |
| --- | --- | --- | --- | --- | --- |
| *FU841_RS06475* | C05625 | Rutin | -1.0000 | 0.0000 | *pepB*; aminopeptidase PepB |
| *FU841_RS06475* | C05903 | Kaempferol | -0.9940 | 0.0060 | *pepB*; aminopeptidase PepB |
| *FU841_RS06475* | C00389 | Quercetin | -0.9800 | 0.0200 | *pepB*; aminopeptidase PepB |
| *FU841_RS13530* | C01477 | Apigenin | -0.9960 | 0.0040 | *ydfG*; bifunctional NADP-dependent 3-hydroxy acid dehydrogenase/3-hydroxypropionate dehydrogenase YdfG |
| *FU841_RS13530* | C01514 | Luteolin | -0.9930 | 0.0070 | *ydfG*; bifunctional NADP-dependent 3-hydroxy acid dehydrogenase/3-hydroxypropionate dehydrogenase YdfG |
| *FU841_RS08345* | C00814 | Biochanin A | -0.9780 | 0.0220 | adenine deaminase |
| *FU841_RS14145* | C05903 | Kaempferol | -0.9740 | 0.0260 | *paaJ*; phenylacetate-CoA oxygenase subunit PaaJ |
| *FU841_RS14145* | C00389 | Quercetin | -0.9750 | 0.0250 | *paaJ*; phenylacetate-CoA oxygenase subunit PaaJ |
| *FU841_RS15025* | C00389 | Quercetin | -0.9750 | 0.0250 | *kduD*; 2-dehydro-3-deoxy-D-gluconate 5-dehydrogenase KduD |
| *FU841_RS15025* | C05903 | Kaempferol | -0.9740 | 0.0260 | *kduD*; 2-dehydro-3-deoxy-D-gluconate 5-dehydrogenase KduD |
| *FU841_RS07090* | C01514 | Luteolin | 0.9980 | 0.0020 | *eutG*; ethanolamine utilization ethanol dehydrogenase EutG |
| *FU841_RS07090* | C01477 | Apigenin | 1.0000 | 0.0000 | *eutG*; ethanolamine utilization ethanol dehydrogenase EutG |
| *FU841_RS07105* | C01514 | Luteolin | 0.9980 | 0.0020 | *eutB*; ethanolamine ammonia-lyase subunit alpha |
| *FU841_RS07110* | C01477 | Apigenin | 0.9990 | 0.0010 | *eutC*; ethanolamine ammonia-lyase subunit EutC |
| *FU841_RS12940* | C05625 | Rutin | 0.9980 | 0.0020 | NAD(P)-dependent oxidoreductase |
| *FU841_RS08200* | C01477 | Apigenin | 0.9990 | 0.0010 | *cdd*; cytidine deaminase |
| *FU841_RS08200* | C01514 | Luteolin | 1.0000 | 0.0000 | *cdd*; cytidine deaminase |
| *FU841_RS14155* | C01514 | Luteolin | 0.9990 | 0.0010 | *paaB*; 1,2-phenylacetyl-CoA epoxidase subunit B |
| *FU841_RS14155* | C01477 | Apigenin | 1.0000 | 0.0000 | *paaB*; 1,2-phenylacetyl-CoA epoxidase subunit B |
| *FU841_RS07080* | C01477 | Apigenin | 1.0000 | 0.0000 | aldehyde dehydrogenase EutE |

Together with Vitamin B2, L-threonine was also found to be negatively correlated with 2 ABC transporters genes (Table 3). Termed as *livF, FU841_RS01625* was one of the open reading frames in a gene cluster, that encodes putative membrane-associated protein [37]. Another gene, *FU841_RS19145* was annotated as a sugar ABC transporter ATP-binding protein related one. It was well known that Gram-negative bacteria are surrounded both by inner and outer membrane which serve as a protective barrier to limit entry of antibiotics. The ATP-binding cassette (ABC) transporter was located in the inner membrane that powers transport [38]. These results suggest that Rutin induced the accumulation of Vitamin B2 and L-threonine and inhibited the expression of ABC transporter related genes, further blocked the activity of ABC transporter.

As a bacterial enzyme, Ethanolamine ammonia-lyase (EAL) catalyzes the adenosylcobalamin-dependent conversion of certain vicinal amino alcohols to oxo compounds and ammonia [39]. Ethanolamine had been proved to promote bacterial growth [40, 41]. The *eut* operon consists of 17 genes [42] and encodes proteins essential for the cobalamin-dependent utilization of ethanolamine [43]. Among those genes, *eutA* encodes a reactivating factor of EAL [44], *eutB* and *eutC* encode the large and small subunits of EAL, respectively [39]. Besides, *eutE* or *eutG* exhibited ability of utilizing ethanolamine as a carbon source and nitrogen source [45]. In present study, L-threonine, 2,3-Dihydroxybenzoic acid and 3 flavonoids (Quercetin, Naringenin and Kaempferol) were found to be positively correlated with EAL related genes (*eutA, eutC* and *eutG*). For example, *FU841_RS07110* (*eutC*) was up regulated to more than 6 fold. Therefore, we hypothesized that the up regulation of a multitude of *eut* genes enhanced the utilization rate of ethanolamine, while the depletion of ethanolamine constrained bacterial growth. However, more evidence is still needed to prove this assumption.

Similarly, the differences between the two treatments, as presented in Table 4, may potentially contributed to the disparate effects observed for Rutin and Luteolin. Rutin, together with Kaempferol and Quercetin showed negative correlation with *pepB* (aminopeptidase PepB). It has been proved that protein degradation by aminopeptidases is involved in bacterial responses to stress [46]. The accumulation of two flavonoids, Quercetin and Kaempferol was found to be negatively correlated with *kduD* gene (Table 4). This gene encodes the KduD enzyme which catalyzes the reduction of 5-keto-D-gluconate, the oxidation of D-gluconate and the conversion of novel sugar substrates [47]. This enzyme has the ability for oxidation but the flavonoids could be used as antioxidants. As a result, it can be concluded that flavonoids possibly inhibited the activity of KduD enzyme and limited these biological process.

### 3.6 Gene validation

The general effect of Rutin is to enhance metabolites production while suppressing genes’ expression. From qRT-PCR results, most of the genes were down regulated after Rutin treatment (Fig. 6 A, B). However, *ompk26* gene was not significant in wild control group and experimental group. All the other genes showed significance between control and experimental groups. Further analysis showed that the expression level of *ompk26* gene was very low, this could be the reason that no significance was found. Similarly, another gene *FU841_RS04100* gene was also weakly expressed and showed no significance between wild control and treatments.

**Fig. 6.**
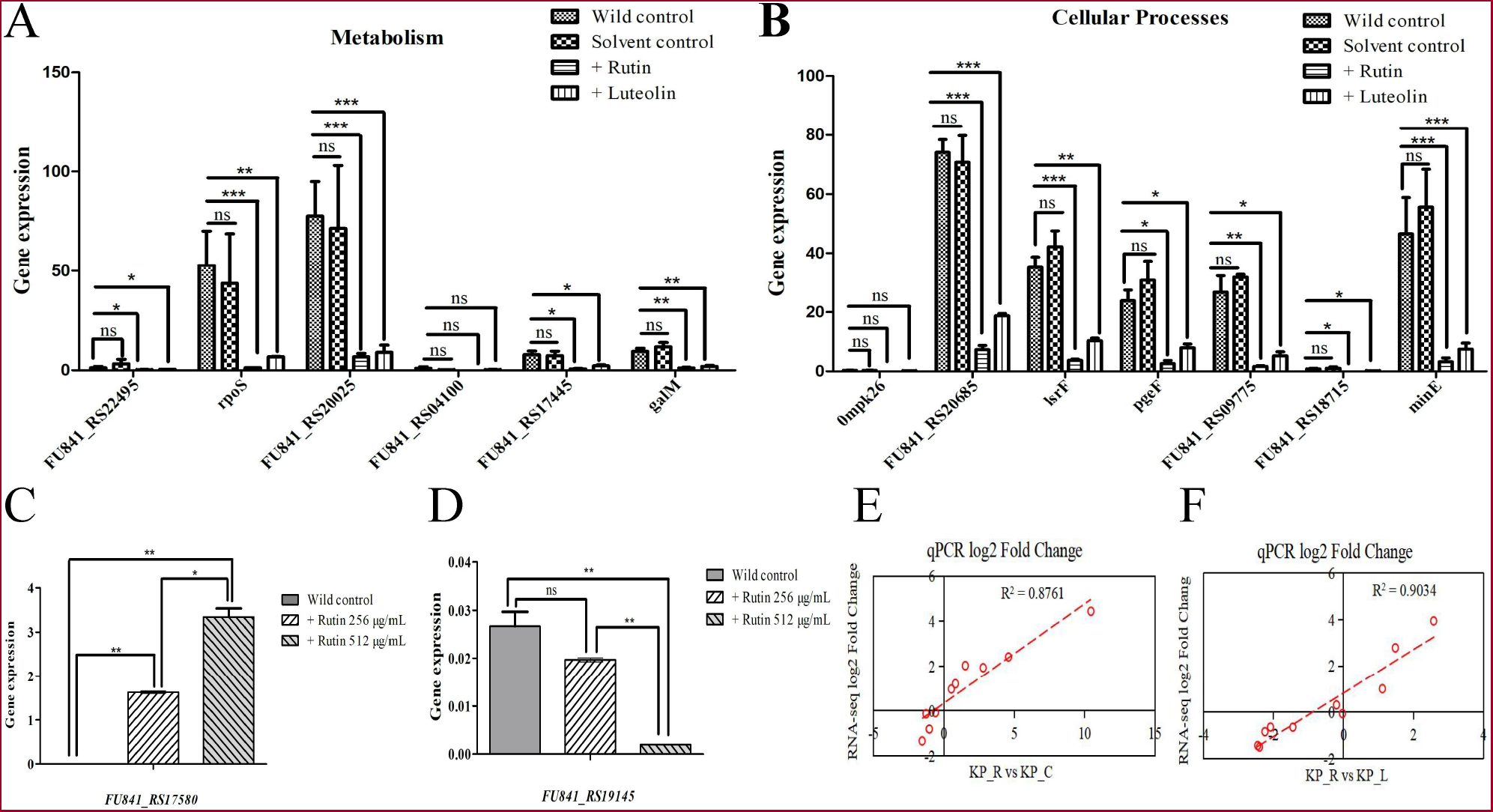
Relative expression profiles and validation of DEGs. (A) Metabolism genes’ expression after flavonoid treatments for 4 hours. (B) Cellular processes genes’ expression after flavonoid treatments for 4 hours. (C)The expression of *FU841_RS17580* after Rutin treatment. (D) The expression of *FU841_RS19145* after Rutin treatment. (E) Correlation of 10 selected DEGs gene expression (log2FC) between RNA-seq and qRT-PCR validation in two pairwise comparisons.* indicates P<0.05, ** indicates P<0.01 and ns indicates not significant.

Gene *FU841_RS17580* might be specifically induced by Rutin as no transcript was obtained in wild control from RNA-seq data. However, low expression of *FU841_RS17580* was measured in wild control by qRT-PCR (Fig. 6 C). Furthermore, when the concentration of Rutin was 256 μg/mL, the expression of *FU841_RS17580* was up regulated. When Rutin’s concentration increased to 512 μg/mL, the relative expression level of *FU841_RS17580* was higher. Gene *FU841_RS17580* was responsible for malonyl-ACP O-methyltransferase. Previous studies expounded the tight correlation between *bioC* and biotin, the latter plays important roles in bacteria growth.

Another gene, *FU841_RS19145* was a ABC transporter related gene and found to be suppressed by Rutin treatment (Fig. 6 D). In qRT-PCR, when the concentration of Rutin increased, *FU841_RS19145* decreased its expression. The highest expression level was detected in wild control while the lowest expression was found in the group which was treated by 512 μg/mL Rutin. These results indicated that the expression of ABC transporter related gene was suppressed by Rutin and the activity of intracellular transport activity was suppressed.

To validate the results of RNA-seq, 10 candidate genes were randomly selected for qRT-PCR and their relative expression was analyzed. Results showed that the expression levels of genes detected by RNA-seq were consistent with that of qRT-PCR method (Fig. 6 E, F). The primers used in this study were listed in Supplementary material, Data_sheet_1, Table S 8.

## Discussion

The emergence of antimicrobial resistance brings great challenges to clinical anti-infective therapy. This underlines the urgent need for new discovery and development of effective drugs [48]. Flavonoids are increasingly attracting attention among researchers not only because they had been indicated to possess effective properties against pathogenic microorganisms [49], but also because these phytomolecules are free of toxicity and environmental hazards [50]. However, a series of studies have focused on the inhibitory effects of flavonoids against bacteria but gained no great success [51]. Based on our previous studies, we found the flavonoids had the characteristics of wide range of property but weak function. Therefore, it is important to reveal the different work mechanism of flavonoids.

Although flavonoids have been reported to possess a wide range of pharmacological properties, like antimicrobial and antiviral activities [52], the antibacterial activities of different flavonoids components varied [53]. Rutin, a kind of flavonols, showed efficient biofilm inhibition of *Streptococcus suis* without impairing its growth [54]. Quercetin was reported not only to inhibit the carbapenemase and efflux pump, but also exerted bactericidal activity through disrupting cell-wall/membrane integrity and altering cellular morphology when used in combination with meropenem in carbapenem-resistant *K. pneumoniae* strain [55, 56]. Another study demonstrated that Rutin showed higher antibacterial activity against *K. pneumoniae* RSKK 574 standard strain than its ESBL+ isolates [57]. According to our previous studies, Rutin was found to exhibit growth inhibition against *K. pneumoniae* strains, but other flavonoids showed weak or even no inhibitory effects [27]. In this regard, we hypothesized the inhibitory mechanism of Rutin is unique and different to other flavonoids.

Rutin is a flavonol compound and luteolin is a flavone. Although both of them have four hydroxyl, there are still three different substituents (R1, R5 and R7). In present study, 12 and 4 pathways were identified from the KP_R vs KP_C and KP_R vs KP_L group, respectively. As listed in Table 3 And 4, only the *eut* genes cluster were similar, the other genes were different. It indicates that the activity of Rutin was different to that of Luteolin possibly due to their different structures. This study demonstrated that Rutin possibly showed inhibition through inducing changes in biotin biosynthesis, ethanolamine metabolism and ABC transporter system, further influenced the strain growth. However, Luteolin was showed to be involved in other biological processes besides the ethanolamine metabolism.

Ethanolamine metabolism plays vital role in strain growth. EAL or EAL related genes were identified in both Table 3 And Table 4. These genes encode proteins that are essential for the cobalamin-dependent utilization of ethanolamine [43]. All the DAMs correlated with EAL or EAL related genes showed positive correlation. Herein, the accumulation of DAMs induced by Rutin promoted the expression of those EAL genes, further leaded to the consumption of ethanolamine. It had been reported that ethanolamine could promote bacterial growth [40, 41]. As a result, the lack of ethanolamine could be a inhibitory factor that blocked the strain growth.

In present study, we measured 5,483 genes and 882 metabolites and identified DEGs and DAMs from both the transcriptomic and metabolomic sequencing data. However, some of the P value in enrichment analysis was greater than 0.05. Therefore, further validation of these results will be our future work. We uncovered that the inhibition of Rutin against *K. pneumoniae* strains mainly due to the its modulation of the metabolic pathways and ABC transporters pathways. Although Rutin could specifically induce the expression of *FU841_RS17580* and up regulate malonyl-ACP O-methyltransferase, we can not draw the conclusion that a single gene or metabolite could inhibit the strain growth. Based on the network, we speculated that Rutin could limit the expression of *FU841_RS19145* and further inhibit the activity of ATP-binding protein, then suppress strain growth.

From the metabolomics, 13 DAMs were identified in both comparative groups. In addition to 4 DAMs that they shared, two comparative groups also have their own unique metabolites. The unique substances identified in KP_R vs KP_C were L-threonine, Naringenin, Vitamin_B2, C00196 (2,3-Dihydroxybenzoic_acid, C7H6O4) and Phenylacetaldehyde. Each of these metabolites was correlated with several genes. Among the 4 kinds of metabolites, Apigenin, Luteolin and Formononetin were flavonoids, and one was Guanosine. It could be supposed that different flavonoids could induce different changes in strain cells. Some of the changes could block strain growth while some others could not.

*Klebsiella pneumoniae* is a significant pathogenic factor of severe infections in humans. Due to its natural resistance to antibiotics, infection with this pathogen can cause severe therapeutic problems. However, with the emergence of drug-resistant strains, it is difficult to choose the right type of conventional antibiotics in clinical practice. In present study we provided valuable information about the response of *K. pneumoniae* to flavonoid treatments. The two flavonoids have both the same and different effects. Vitamin B2 and L-threonine were specifically induced after Rutin treatment and showed negative correlation with ABC transporter related genes. Our results revealed the potential inhibitory mechanism of Rutin on strain growth as Rutin induced the accumulation of Vitamin B2 and L-threonine, and also inhibited the expression of ABC transporter related genes, further suppressed strain growth. However, further study is still needed to learn how different flavonoids influence the growth of *K. pneumoniae*, especially to those clinical isolates.

## Conclusions

The present study suggested that Rutin might induce changes in biotin biosynthesis, ethanolamine metabolism and ABC transporter system and further suppress the strain growth. In addition, the inhibitory mechanism of Rutin was different to that of Luteolin. These findings derived from integrated multi-omics study and intracellular experiments are encouraging for new development of antimicrobial agents and pathogenic microbial infection control.

## Supplementary Information

The online version contains supplementary material.

## Acknowledgments

We appreciate the funding support from Luzhou Government and Southwest Medical University Cooperation Program (2020LZXNYDJ46).

## Authors’ Contributions

The authors ZW, WS and XW conceptualized the study. ZW, YL, X-YW, GX and HT performed the experimental work and collected the data. ZW, NM and XY analyzed the data and prepared the manuscript. ZW and WX performed all bioinformatic analyses and wrote the manuscript. WX and XW reviewed the manuscript. XW and XZ supervised the project, experimental design, the data collection and analysis, and manuscript preparation. All authors have read and approved the manuscript for publication.

## Availability of data and materials

The RNA-seq datasets generated and/or analysed during the current study are available in the NCBI GenBank (https://www.ncbi.nlm.nih.gov/genbank/), the accession number of our submission is BioProject: PRJNA956990 or BioSample: SAMN34237808. The raw data of the metabolomics was deposited in figshare (https://figshare.com/) and published (DOI:10.6084/m9.figshare.22884191).

## Declarations

The authors claim that none of the material in the paper has been published or is under consideration for publication elsewhere. The submission is original, and all authors are aware of the submission and agree to its publication in Microbial Pathogenesis.

## Ethics approval and consent to participate

Not applicable.

## Consent for publication

Not applicable.

